# Correlated Light-Serial Scanning Electron Microscopy (CoLSSEM) for ultrastructural visualization of single neurons *in vivo*

**DOI:** 10.1101/148585

**Authors:** Yusuke Hirabayashi, Juan Carlos Tapia, Franck Polleux

## Abstract

A challenging aspect of neuroscience revolves around mapping the synaptic connections within neural circuits (connectomics) over scales spanning several orders of magnitude (nanometers to meters). Despite significant improvements in serial section electron microscopy (SSEM) technologies, several major roadblocks have impaired its general applicability to mammalian neural circuits. In the present study, we introduce a new approach that circumvents these roadblocks by adapting a genetically-encoded ascorbate peroxidase (APEX2) as a fusion protein to a membrane-targeted fluorescent reporter (CAAX-Venus), and introduce it in single pyramidal neurons *in vivo* using extremely sparse *in utero* cortical electroporation (IUCE). This approach allows to perform Correlated Light-SSEM (CoLSSEM) on individual neurons, reconstructing their dendritic and axonal arborization in a targeted way via combination of high-resolution confocal microscopy, and subsequently imaging of its ultrastuctural features and synaptic connections with the ATUM-SEM (automated tape-collecting ultramicrotome - scanning electron microscopy) technology. Our method significantly improves the the feasibility of large-scale reconstructions of neurons within a circuit, and bridges the description of ultrastructural features of genetically-identified neurons with their functional and/or structural connectivity, one of the main goal of connectomics.

---

Unbiased, saturated connectomic approaches are aimed at reconstructing all synaptic connections within neural circuits at ultrastructural resolution [1-3]. Despite significant technological advances, current approaches cannot be generally applied to the description of large-scale neuronal connectivity at ultrastructural resolutions because of the following major roadblocks[4]. Current SSEM approaches are mainly applied to small pieces of brain tissue (~hundreds of cubic microns), which is incompatible with the mapping of long-range connections organized over millimeters to meters[5]. Within a given volume, the identity of each of the elements in the circuit (cell types, axons, dendrites, synapses) is undetermined and there are limited ways to identify the axons or dendrites originating outside the volume reconstructed by SSEM.

Fluorescent light microscopy (LM) however, coupled to genetic labeling of neurons, permits the identification and tracking of axons, dendrites and their branches over long distances in intact behaving animals. Unfortunately, this approach only informs about target regions, giving no information about synaptic partners. Therefore, the correlation of LM to electron microscopy-imaged structures (CLEM) could overcome the problems associated to the study of long range circuits[6]. Although many efforts have been performed in this direction, a prevailing limitation of most CLEM studies is that the tools used to label specific cells obscure most of their ultrastructural features severely affecting their implementation. In the meantime, there is an urgent need to characterize single neurons at the molecular, cellular and circuit levels. Because individual synapses have dimensions below the light diffraction limit, neuronal circuit reconstructions at ultrastructural levels require the use of either super-resolution microscopy or serial electron microscopy (SEM). The advantage of EM approaches over super-resolution microscopy is that it provides rich and unbiased ultrastructural information of synapses as well as details about the organelles and subcellular elements that compose individual neurons[7, 8]. Several recent studies have attempted to correlate the functional and structural properties of neuronal ensembles defined by fluorescent imaging with the mapping of their synaptic connections using serial EM [9, 10]. However, several roadblocks limit the general applicability of such approaches to the mammalian central nervous system (CNS): (1) the difficulty of mapping long range connections (hundreds of microns to millimeters-meters) between genetically- and/or functionally-identified neurons within a circuit, and (2) the fact that most genetically-encoded EM contrasting agents (HRP, APEX, miniSOG) diffuse in the cytoplasm and obscure the neurons ultrastructural information disabling a detailed description of their subcellular features[11, 12].

A recently developed monomeric peroxidase reporter, APEX2[13], which keeps its enzymatic activity even in the reducing cytosolic environment of cells or upon fixation, has been shown to be useful for studying protein localization in cultured cell lines and in zebrafish using electron microscopy[14, 15], but its application on mammalian tissue including the brain has not been extensively explored yet. Here, we took advantage of ATUM-SSEM[7] to interrogate at the ultrastructural level large pieces of tissues (~1-4mm) and correlate LM and EM datasets obtained from single mouse cortical pyramidal neurons. We demonstrate that by expressing a plasma membrane-targeted CAAX-Venus-APEX2 fusion protein, we significantly increase the accuracy and feasibility of performing CLEM *in vivo*, allowing the imaging of single layer 2/3 pyramidal neurons and their synaptic connections while preserving the ability to visualize ultrastructural details.

## RESULTS

In order to improve our ability to perform CLEM studies in mammalian neurons *in vivo*, we decided to design a cDNA that encodes a fusion protein between the fluorescent protein Venus and the recently engineered plant ascorbate peroxidase (APEX2;[13]) that catalyzes the polymerization and local deposition of diaminobenzidine (DAB) in presence of H_2_O_2_, which in turn recruits electron-dense osmium responsible for the EM image contrast (Fig. 1A). Since labeling the cytoplasm dramatically interferes with proper visualization of subcellular structures (mitochondria, ER, synapses), a CAAX motif derived from H-Ras was added to the APEX2-Venus genes that tethers the APEX2-Venus-CAAX (AVC) fusion protein to the cell’s plasma membrane (Figure 1A)[16]. We tested the feasibility of the construct primarily in HeLa cells, which after being transfected for 24 hours showed the expected results (Venus fluorescence and AVC-DAB labeling). No phenotypic alterations were observed in these cells (data not shown).

**Figure 1.**
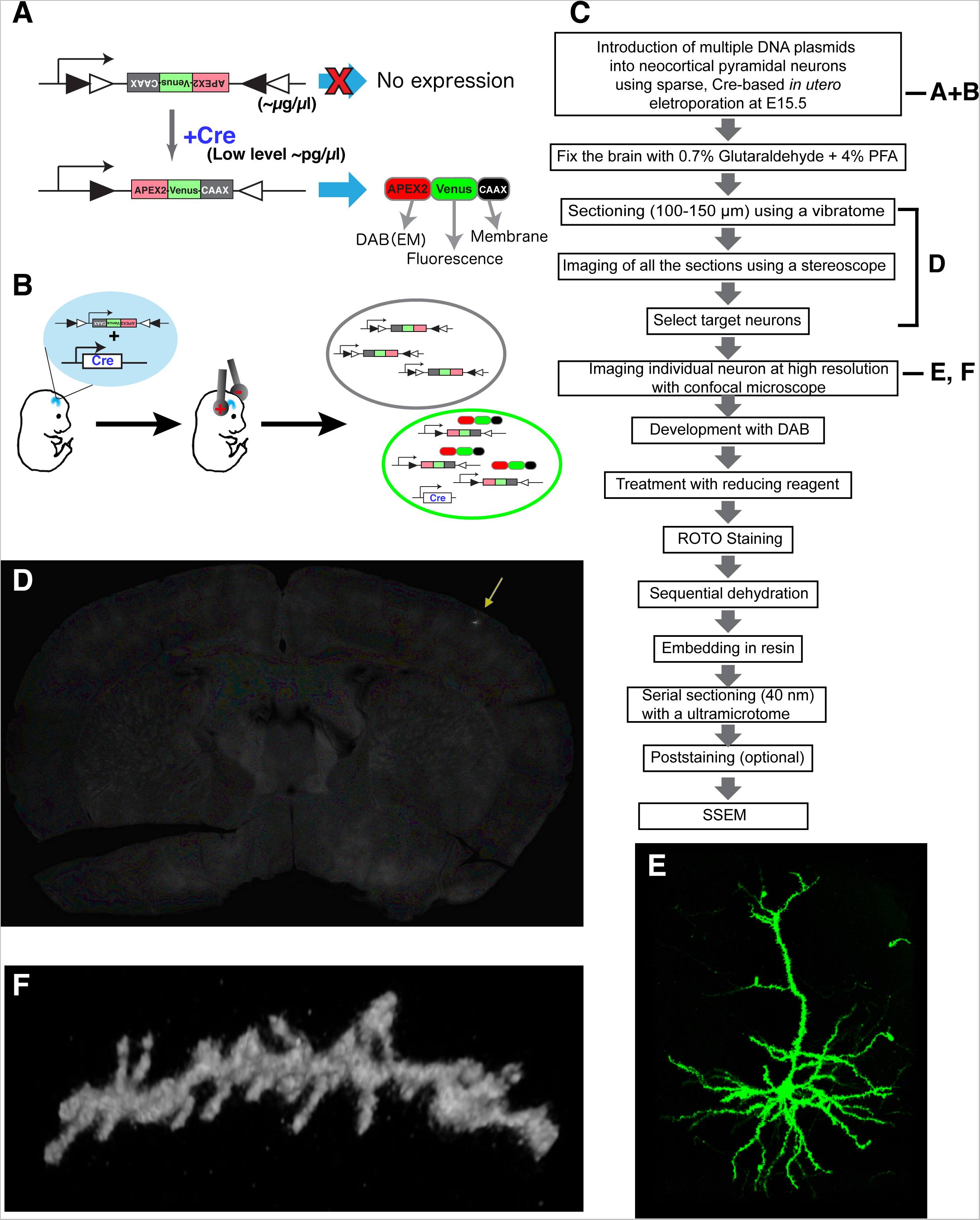
Sparse labeling and fluorescent imaging of individual cortical pyramidal neurons using CAAX-Venus-Apex2 for CoLSSEM. (**A**) Our labeling strategy consists in sparse labeling of single neurons with a Flex plasmid (inverted cDNA flanked by incompatible LoxP sites) which, upon Cre-mediated recombination leads to expression of a fusion protein, APEX2-Venus-CAAX (AVC). DAB, diaminobenzidine. EM, electron microscopy. (**B**) Sparse labeling of individual neocortical pyramidal neurons by *in utero* electroporation. Black circle indicates cells taking up only Flex plasmids. Green circle indicates a cell taking up both Flex and Cre plasmids and expressing APEX2-Venus-CAAX. (**C**) Flow diagram showing the steps involved in CoLSSEM. (**D**) 100 μm thick coronal section of a brain sparsely labeled with Venus (arrow) at postnatal (P) 25. (**E,F**) 3D reconstruction from confocal images of a layer 2 cortical neuron at P25 (E, 74 z-stack images with 0.9 μm z-step) and a dendritic segment with spines at P23 (F, maximum projection of 17 z-stack images with 0.8 μm z-step) labeled with AVC fusion protein (also see Supplemental movie 2 for 3D animation).

Although genomic integration of marker genes driven by cell-type specific promoters is a versatile and reliable technique for labeling neurons *in vivo*, the dense labeling of cells in small volume significantly hampers the identification of dendritic and axonal processes from individual neurons. To reduce the number of neurons expressing the AVC fusion protein, without compromising its expression level, we decided to build a plasmid that contained an expression cassette flanked by pairs of incompatible LoxP recombination sites (Flex)-system (Fig. 1A)[17]. The Flex plasmid encoding the AVC gene, and a plasmid encoding the Cre recombinase gene were mixed at 10,000:1 dilution ratio, and injected into the lateral ventricle of a mouse embryo (E15.5). The plasmids were introduced into neural progenitor cells in the ventricular zone by using *in utero* cortical electroporation[18-20]. Using such low concentration of Cre-expressing plasmid (~100-200 pg/μl) leads to stochastic recombination of the Flex-AVC plasmid in progenitors of layer 2-3 pyramidal neurons in the ventricular zone of the dorsal telencephalon leading to very sparse expression (Fig. 1B)[21].

Following long-term survival after IUCE, mouse brains were processed for the CLEM approach (Fig. 1C, Methods). Coronal brain slices were obtained using vibratome sectioning (100 microns), and individual optically isolated Venus-expressing pyramidal neurons in layer 2-3 were identified by imaging all serial vibratome sections from single adult mouse brains, covering the entire somatosensory area, at low magnification using a fluorescent stereoscope (equipped with a X-Y motorized stage; see Methods) (Fig. 1D). Using this approach, the number of neurons expressing the AVC fusion protein varied among individual mouse brain from 10-30 neurons/brain. These optically isolated pyramidal neurons were then imaged using a confocal microscope, and high resolution image stacks were collected to document the detailed dendritic and axonal morphology of each labeled neuron at sub-micron resolution (Figure 1E-F, see also example in Figures 2A and 4A, **and Supplemental Movies S1, S2 and S4)**. Analysis of the spine density in apical oblique dendrites of layer 2/3 neurons at P23 (1.05 ± 0.12 spines/μm, mean ± s.d.) as well as the number of primary dendrites indicates that AVC fusion protein is innocuous for these neurons. Moreover, this LM approach provided information essential for determining the morphological features of individual layer 2/3 pyramidal neurons as shown in Figure 1, and to track their long-range axonal projections (see Figure 4).

**Figure 2.**
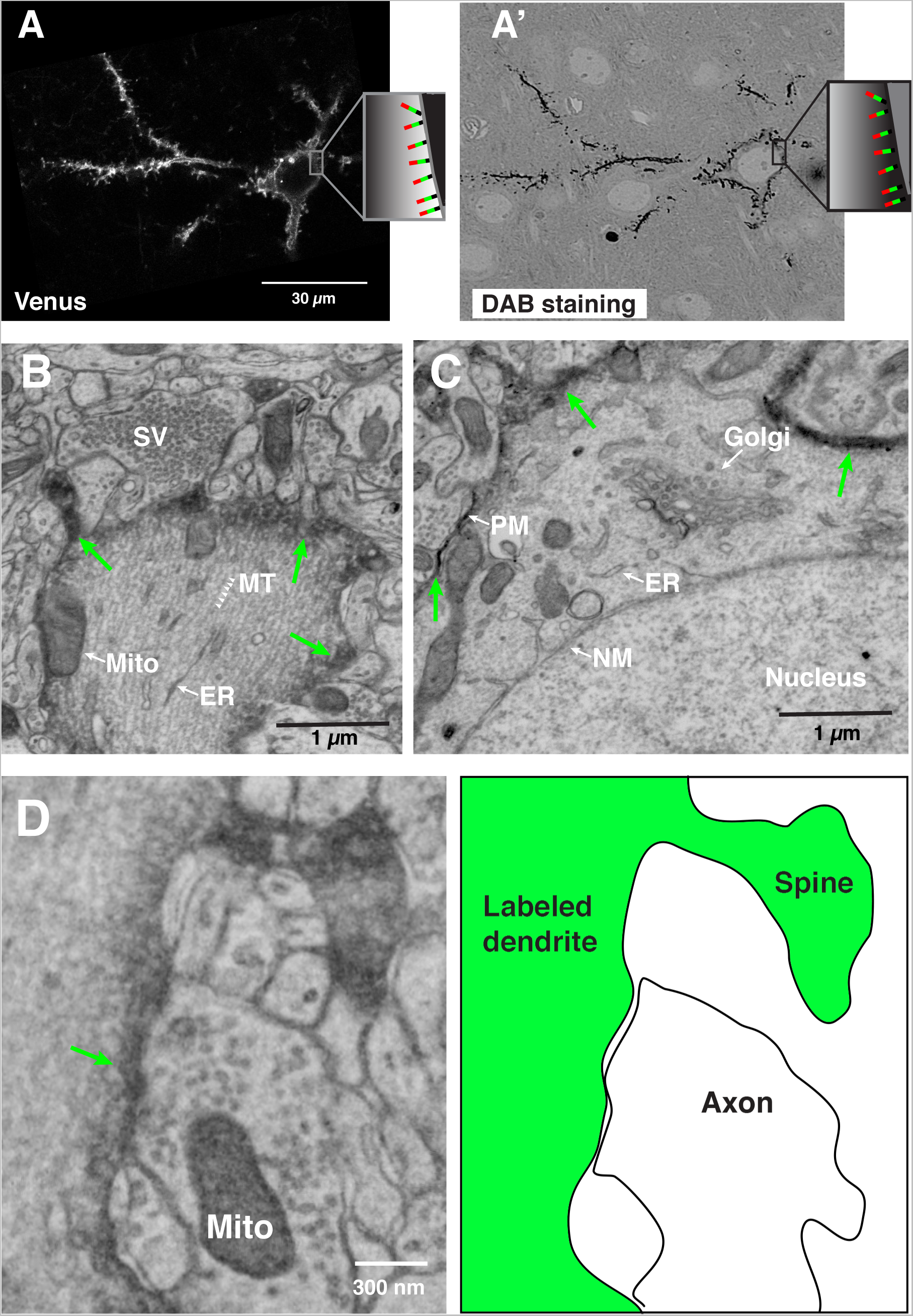
Correlative LM-EM labeling of single neocortical neurons. (**A-A’**) Correlated images of Venus imaged by confocal microscopy (A) and DAB signal taken by the electron microscope (A’) from a single neuron labeled with APEX2-Venus-CAAX protein. (Inset; Schema illustrating the topology of the AVC protein on the plasma membrane; Red, Apex2. Green, Venus. Black, CAAX motif). (**B-D**) Single scanning electron micrographs showing the intracellular structures of a single labeled pyramidal neuron in layer II/III at P25. Green arrows indicate the plasma membrane labeled with APEX2-mediated DAB staining. (**B**) Primary apical dendrite. Arrowheads indicate microtubules (MT). (**C**) Cell body and its resident organelles: the Golgi apparatus, Nuclear membrane (NM), and the plasma membrane (PM) labeled with electron-dense DAB from APEX2. (**D**) Electron micrograph showing a primary dendrite, a spine, and a presynaptic terminal targeting to the dendritic shaft (Left). Diagram illustrating the structures shown in the electron micrograph (right). Abbreviations: SV, synaptic vesicle. MT, microtubule. Mito, mitochondria. ER, endoplasmic reticulum.

Following treatment with DAB and H_2_O_2_, the membrane targeted APEX2 converted the fluorescently tagged neurons (CAAX-Venus) into a dark, electron-dense precipitate (Fig. 2A-A’), compatible with EM detection (discussed below). Sections containing the labeled neurons were processed for SSEM as previously described[7, 22]. To align the LM and EM images in the z-dimension, we placed each section between two plastic plates and kept the sections flat during the resin embedding process (see Methods). Then resin-embedded thin sections were collected by ATUM in the same plan as the confocal images. The neurons imaged under the LM were re-identified in the EM images by visual inspection allowing correlated light-EM microscopy (Figure 2A-A’). Whereas the subcellular membrane structures were partially disrupted in the images obtained from the brain fixed with 0.25% glutaraldehyde (GA) and 4% paraformaldehyde (PFA) (data not shown), they were well preserved and DAB staining was optimal when GA concentration was increased to 0.7% (Figure 2 B-D). The DAB staining at the plasma membrane clearly labels the target neuron, but importantly does not interfere with ultrastructural details of organelles, cytoplasmic content and nuclear structures (Figure 2 B-D).

To assess the consistency of the DAB-AVC mediated staining, and examine the detailed ultrastructure of the AVC-labeled neuron, we performed a reconstruction of a segment of the labeled dendrite from the SSEM images. Generally, small molecule penetration into glutaraldehyde fixed tissue can be a limiting factor[23]. Consistent with this idea, simultaneous incubation of tissue sections with both H_2_O_2_ and DAB substrates for 5-15 mins did not give us a clear, well defined staining. However, a pretreatment of the glutaraldehyde fixed vibratome sections with the DAB substrate for 30 minutes, prior the treatment with H_2_O_2_, significantly improved the DAB-AVC mediated deposition, providing evenly stained neuronal processes throughout the 100 μm section (Figure 3A, a summary of the optimization Figure 3D). Then, we reconstructed the apical dendritic trunk from DAB-labeled layer 2/3 neurons by manual tracing (Figure 3B). Despite the strong staining following DAB pre-incubation, all cytoplasmic organelles such as mitochondria, endoplasmic reticulum (ER) and Golgi apparatus were easily identified (Figure 3 A,C **and Supplemental Movie 3)**.

**Figure 3.**
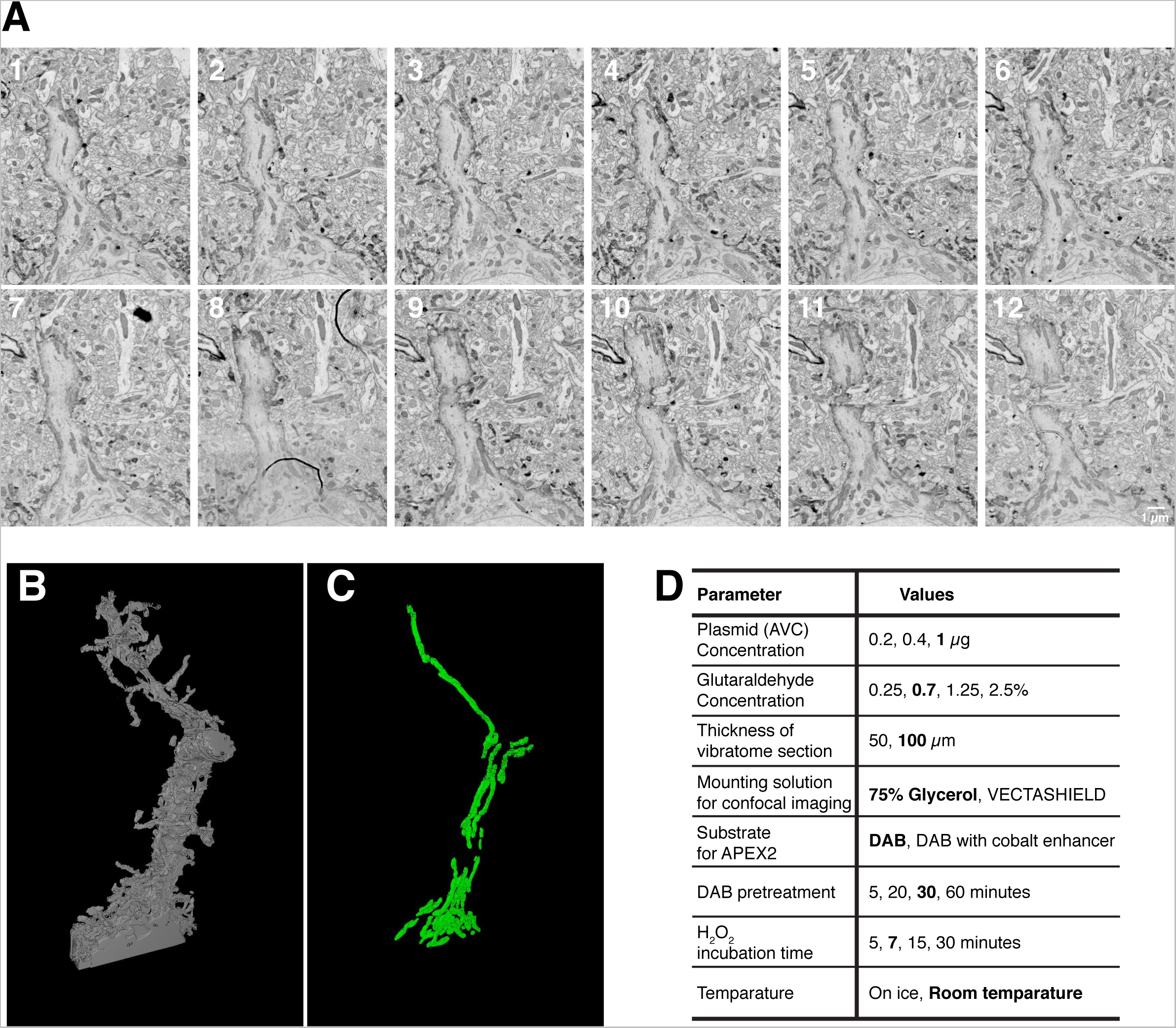
3D-SEM reconstruction of the primary dendrite of single AVC-labeled pyramidal neuron and its subcellular structures. (**A**) Montages of serial electron micrographs including the DAB labeled pyramidal neuron shown in Figure 1d-e. (see Supplemental movie 3). (**B,C**) 3D reconstruction of the plasma membrane (B) and the mitochondria (C) of its labeled apical dendrite (see Supplemental movie 4 and 5). To find out the region of interest, 256 serial sections were imaged with 30 nm/px resolution. The region of interest was re-imaged with 3 nm/px resolution through 103 serial sections and reconstructed. (**D**) Parameters tested to optimize the DAB staining and preservation of the membrane structures for SSEM. Bold indicates the optimal conditions used in the paper.

Tracing axonal segments over long distances from genetically-identified neurons would be a major advance for connectomics studies. However, it has been challenging to perform axonal reconstructions from SSEM images because axons are long (>hundreds of microns) and thin (<1 micron diameter), adopting tortuous 3 dimensional trajectories. By using DAB-AVC method, since the axon of interest is well labeled by the DAB staining protocol, compared to non-genetically transfected processes, and confirmed by the high resolution fluorescent imaging, we were able to trace the axon on SSEM images further away from its cell body. As a result, we could perform correlated light- and SSEM imaging of individual axons, and unambiguously trace a single axon and its branches for >200 μm long or more in the EM images (Figure 4 A-C **Supplemental Movies 4-7)**.

**Figure 4.**
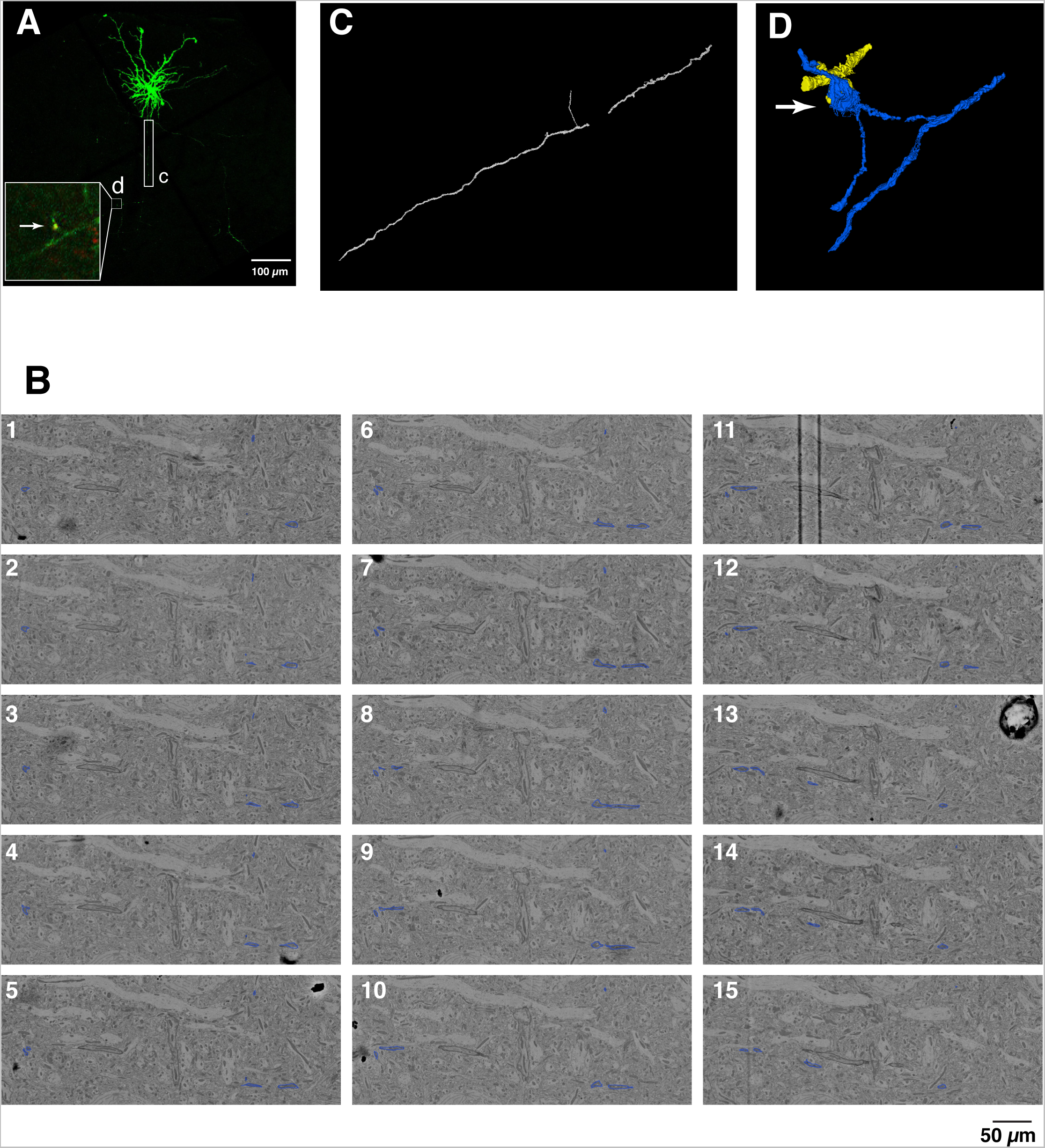
3D correlative imaging of neocortical axons. Confocal images and electron micrographs of a neuron (P28) electroporated with AVC together with vGlut1-mCherry as a presynaptic marker. (**A**) A maximum projection image of a layer 2/3 neuron projecting to layer 5. (Inset) Close-up view of the area indicated by a square, which is reconstructed by SSEM in (D). The presynaptic site is labeled with vGlut1-mCherry (red-inset). Rectangle, the axonal part reconstructed in (C). (**B**) Montages of electron micrographs including the single DAB labeled axon (see Supplemental movie 6 and 7). The DAB labeled axon was manually segmented (blue). (**C**) 3D reconstruction of the axon indicated by a rectangle in (A). 216 serial sections were imaged with 20 nm/px resolution. The region of interest was re-imaged with 8 nm/px resolution through 103 serial sections and reconstructed. (**D**) 3D reconstruction of the labeled axon and presynaptic site identified by vGlut1-mCherry labeling (blue), and its postsynaptic target (yellow).

The challenge represented by tracing a single axon of a genetically-identified neuron for a long distance is a major roadblock for identifying presynaptic boutons of a specific neuronal subtype in the SEM images. The correlated imaging of Venus and DAB circumvents this limitation by re-identifying individual presynaptic boutons identified in the LM images. The presynaptic bouton of layer 2/3 pyramidal neuron in layer 5 was identified by the enrichment of vGlut1-mCherry on the Venus labeled axon in LM (Figure 4A). By correlating this site in the EM sections using DAB staining, the structure of this presynaptic site and its apposed postsynaptic structure were reconstructed from SSEM imaging (Figure 4D **and Supplemental Movie 8).**

## Discussion

In this study, we combined a genetic approach to label a single neuron with a novel fusion protein allowing to perform correlated light- and SSEM imaging (CoLSSEM). The use of two image modalities (LM and EM) allows for the first time true examination of ultrastructural features of individual genetically-identified neurons as well as identification of synaptic connection of a specific type of neuron in mammalian neocortex. These results show that CoLSSEM improves the accuracy and feasibility of linking multiscale neuronal projections i.e., from mm^3^ to nm^3^.

By adopting a recently developed peroxidase APEX2 and targeting it to the plasma membrane as a fusion with Venus (Apex2-Venus-CAAX), we have been able to successfully ‘highlight’ the plasma membrane of single neurons *in vivo*. Since enzymatic activity of APEX2 is preserved after treatment with chemical fixatives such as p-formaldehyde (PFA) and glutaraldehyde, the short-term (5-10 minutes) treatment with H_2_O_2_ in presence of DAB substrate allows optimal staining to highlight a target cell plasma membrane from the surrounding non-labeled cells. Short-term treatments are beneficial, since long term incubations with H_2_O_2_, which is required for the staining with HRP (~1.5 h), alter significantly the integrity of membranes and neuritic processes. Further, LM-EM dual labeling from a single protein enabled unambiguous correlation of the neurons between LM and EM. Increased image quality and reliability of light-SSEM correlative imaging helps precise description of ultrastructural features of single cells in complex tissues such as the brain.

Future application of our new approach will need to test the use of Flex-AVC probe and express it in transgenic mice expressing Cre recombinase in genetically identified cell classes within the developing and adult brain. The rapid development of new mouse lines expressing Cre in a cell-type specific way will allow sparse or dense labeling of ensembles of neurons and perform mapping of the synaptic connectivity within circuits at ultrastructural resolution[24-27].

In addition to the labeling by genetically encoded probes, the combination of the sparse labeling and ATUM-SSEM significantly improves the efficiency for re-identification of the labeled neurons. The genetic approach we adopted in this study labeled neurons sparsely enough to unambiguously trace the projection of individual neurons, which improves correlating LM and EM. The ATUM-SSEM approach enabled us to preselect the regions containing the sparsely labeled neuron efficiently using fast, large-area and low-resolution imaging using a stereoscope equipped with a X-Y motorized stage and a very sensitive CCD camera. In the future, improvement of large-scale, single axon imaging[28], in combination with our CoLSSEM approach will allow targeted, three-dimensional reconstructions of axons and dendrites at nanometer resolution.

Besides the correlation of LM and EM, the sparse labeling of neurons with our new AVC probe will improve connectomic approaches. We traced the axon from layer 2/3 neocortical neuron for 200 μm, which is significantly longer than the tracing performed in a previous study using ATUM-SSEM (50-60 μm; [29]). The membrane contrast provided by our new membrane-targeted APEX2 labeling strategy will also improve automated tracing of neuronal processes based on segmentation, which is a key step for the emergence of large scale connectomic studies[30]. Given that our LM-EM dual labeling technique can be applied to any cell types by LM by fusing APEX2 with an appropriate fluorescent protein, such as genetically encoded calcium indicators, CoLSSEM will improve the description of the ultrastructural features and the connectivity characterizing specific cell types within neural circuits *in vivo*. The ability to combine this strategy with other approaches (*in vivo* morphological and functional imaging) will provide critical new insights into the relationship between neuronal properties, circuit connectivity and its emerging functional properties underlying complex behaviors such memory formation or sensory perception and cognition.

## Legends for supplementary movies

**Movie S1.** Related to Figure 1E. Low magnification, 3D rotating projection of confocal stack of single cortical pyramidal neuron in layer 2/3 expressing CAAX-mVenus-Apex, imaged at P21.

**Movie S2.** Related to Figure 1E. High magnification and high resolution confocal imaging of a single dendritic branch of pyramidal neuron in layer 2/3 showing single dendritic spines.

**Movie S3**. Related to Figure 3. 3D reconstruction of the apical dendrite of a pyramidal neuron imaged by CoLSSEM, 3D SEM shown in Figure 3.

**Movie S4**. Related to Figure 4. Low magnification, 3D rotating projection of confocal stack of single cortical pyramidal neuron in layer 2/3 expressing CAAX-mVenus-Apex, imaged at P21. The axon segment of this neuron is imaged by 3D SEM in Figure 4.

**Movie S5.** Related to Figure 4. Raw image series used for 3D SEM of axon segment reconstructed in Figure 4.

**Movie S6.** Related to Figure 4. Annotated image series used for 3D SEM of axon segment reconstructed in Figure 4. The axon segment from neuron reconstructed in Figure 4 is outlined in blue.

**Movie S7.** Related to Figure 4. 3D animation of the axon segment imaged by CoLSSEM and traced in Figure 4 (also shown in Movies S5&S6).

**Movie S8.** Related to Figure 4A. 3D animation of the axon segment (blue) imaged by CoLSSEM and traced in Figure 4 focusing on the presynaptic terminal labeled with the genetically encoded vGlut1-mCherry (inset of Figure 4A). The post-synaptic partner of this presynaptic bouton is shown in yellow.

## Methods

### Animals

All animals were housed and handled according to protocols approved by the Institutional Animal Care and Use Committee (IACUC) at Columbia University. Time-pregnant CD1 females were purchased from Charles River (Wilmington MA, USA).

### DNA constructs

The pCAG-vGlut1-mCherry and pCAG-Cre vectors were previously descrived[31]. The pcDNA3 APEX2-NES (Addgene plasmid # 49386) and pAAV-Ef1a-DIO eNpHR 3.0-EYFP (Addgene plasmid # 26966) vectors were gifts from Drs. Alice Ting (Stanford University) and Karl Deisseroth (Stanford University), respectively. The flag-APEX2-NES sequence was cloned into the 5’ end of the Venus sequence and the CAAX Motif–AAGCTGAACCCTCCTGATGAGAGTGGCCCCGGCTGCATGAGCTGCAAGTGTGTGCTCTCCTGA (KLNPPDESGPGCMSCKCVLS) was fused to the 3’ end of the Venus cDNA (APEX2-Venus-CAAX, AVC). Then, APEX2-Venus-CAAX (AVC) sequence was exchanged with the eNpHR-EYFP segment of the pAAV-Ef1a-DIO eNpHR 3.0-EYFP vector by AscI-NheI sites (pAAV-Ef1a-DIO-AVC).

### Introduction of DNA into mouse neocortex

The in utero electroporation method was performed as previously described[32]. Briefly, a mixture of the vectors pAAV-EF1a-DIO-AVC (Flex-AVC; 1μg/μl) and pCAG-Cre (recombinase; 100 – 200 pg/μl) was injected into the neocortical lateral horn of embryos obtained from a timed-pregnant CD1 mouse female at E15.5 with Picospritzer III (Parker, Hollis NH, USA). For co-labeling neuronal processes and their presynaptic sites (Figure 4), The pCAG-vGlut1-mcherry (0.3 μg/μl) were added to the mixture of the vectors indicated above. The cDNA mixtures were then electroporated into the neural progenitor cells resided in the ventricular zone by 5 electric pulses (50 ms) of 38 V using Tweezer-type platinum electrode (diameter 3 mm; NEPA GENE) and an electroporation system ECM 830 (BTX, Holliston MA, USA).

### APEX development and light microscopy imaging of brain sections

Following anesthesia, mice were heart perfused with 5 ml of phosphate buffered saline (PBS 0.1M) and subsequently with 40 ml of the fixative solution containing 4% paraformaldehyde (EM grade, EMS) and 0.25-0.7% of Glutaraldehyde (GA, EMS) diluted in PBS. Brains were then dissected out and post-fixed in the same fixative solution for 3 hours. After embedded in low melting agarose (3%, MP Biomedicals), brains were sectioned at 100 μm thickness with a Leica VT1200S vibratome. Sections were placed on glass slides, mounted in PBS, and covered with coverslips. The series of thick sections were first imaged at low resolution (2.1 μm/px, exposure 1 s) with a 1x (0.15 NA) objective on an automated SMZ18 stereoscope (Nikon) equipped with a fast ORCA Flash 4.0 (Hamamatsu Photonics) and a motorized x-y stage (Prior). Following this process, sections containing individually brightly labeled neurons were remounted in 75% glycerol-PBS imaged at higher resolution with either a 20x (0.75 NA) air objective or 60x (1.4 NA) oil objective on a Nikon Ti-e inverted microscope equipped with a Nikon A1 confocal. Sections were washed (PBS for 5 min) and immersed in a PBS containing the APEX substrate (ImmPact^TM^ DAB Chromogen; Vector) for 30 minutes followed by treatment with ImmPact^TM^ diluent (Vector) for 7 minutes. The strongly APEX-developed sections were washed with PBS (3 times) and treated with 10mg/ml the reducing agent Sodium HydroSulfate (Sigma) in PBS for 10 minutes as indicated[15]. Sections were washed 3 times with PBS, and post-fixed with 2.5% GA overnight. To process for electron microscopy, tissue samples were osmicated using the ROTO staining method (reduced osmium tetroxidethiocarbohydrazide (TCH)-osmium). Sections were followed by incubations in ethanol, propylene oxide (EMS) embedded in epon based resin (EMS). Curing of the resin was achieved at 65°C for 2 days. Epon blocks were trimmed with a TrimTool diamond knife (Trim 45; DiATOME) and ultrathin sectioned with an Ultra Diamond Knife (Ultra 35°; DiATOME) in a Leica Ultramicrotome (UC7) and the serial sections collected using ATUM technology[22, 33, 34]. Approximately 1500 1 mm (width), 3 mm (length) 40 nm thick serial sections were obtained and collected by ATUM using carbon coated tape (kapton).

### Serial section electron microscopy

Placed in a silicon wafer, the region of interest (ROI) in the sections was identified by visualizing blood vessel land marks. Electron micrographs at higher resolution (4nm pixel size, 1ms dwell time) were obtained using a Sigma Field Emission Scanning Electron Microscope (7 KeV; Zeiss) using back-scatter signal detector. All image stacks obtained were then digitally processed, aligned and reconstructed in Fiji (NIH) using the TrackEM2 plugin[35].

## Acknowledgements

This work was supported by grants awarded to NIH (NS089456) (FP) and the Roger De Spoelberch Award (FP), JST and PRESTO (YH), FONDECYT1160888 & ALSA (JCT). We thank Dr. Tom Jessell for his generous support during this project. We thank Dr. Tom Maniatis and Charles Zuker for their advice and generous help during the generation of the manuscript.

